# Macromolecular condensation is unlikely to buffer intracellular osmolality

**DOI:** 10.1101/2024.05.24.592450

**Authors:** Alan R. Kay, Zahra Aminzare

**Affiliations:** Dept. Biology University of Iowa, Iowa City, IA 52242; Dept. Mathematics, University of Iowa, Iowa City, IA 52242

## Abstract

The flux of water across biological membrane is determined by gradients in transmembrane osmolality and the water permeability of membranes. Watson *et al*. (2023) have proposed that the reversible formation and disassembly of molecular condensates could act as the primary buffer of cytoplasmic osmolality in the face of changes in extracellular osmolality. In this communication, we show using well-established membrane biophysics, that the water permeability of plasma membranes is likely to overwhelm any cytoplasmic water buffers.

## Introduction

The regulation of cytoplasmic osmolality is of immense importance for the normal functioning of all animal cells (Krogh, 1946; Larsen et al., 2014). Recently Watson *et al*. (2023) proposed that the reversible condensation of macromolecules could serve to buffer changes in cytoplasmic osmolality induced by extracellular changes in osmolality, a claim which may at first blush seem to be feasible. However, we will show that it is inconsistent with what we know about the water permeability of membranes (Boron and Boulpaep, 2016; Weiss, 1996). It is certainly possible that subjecting macromolecules to changes in osmolality could lead to conformational changes that could induce them to bind or release water (Guttman et al., 1995; Parsegian et al., 2000), hence acting as a water buffer (WB). Nevertheless, as we will show, the known water permeability of the plasma membrane will quickly overwhelm any buffering capacity provided by molecular condensates.

Watson *et al*., show that the relationship between macromolecular concentration and solution osmolality can be highly nonlinear *in vitro*. This has been explored extensively by others, and although precisely how this nonlinearity arises is by no means resolved, the phenomenon is not new (Adair, 1925; Neal et al., 1998; Yousef et al., 1998). Consistent with their hypothesis they observed an increase in cytoplasmic condensate formation with a hyperosmotic challenge to cells, while a hypoosmotic change decreased condensation. However, it is their jump from the nonlinearity of osmolality to concluding that macromolecules can serve to buffer the cytoplasmic osmolality that is questionable, since they provide no direct evidence. It is also worth noting that the formation of condensates can be triggered in giant unilamellar vesicles containing polyethylene glycol and dextran by changes in osmolarity (Li et al., 2008).

In their paper Watson *et al*., refer to the *osmotic potential* (−Ψ_*π*_ = *iCRT*, where *C* is the concentration of the solute, *i* the van ‘t Hoff factor, *R* the gas constant, and *T* the absolute temperature) rather than the osmolarity or osmolality of solutions. Since they measure the osmolality of their solutions, which is the number of moles per kilogram of water (Blaustein et al., 2019), we will use this to quantify the osmotic properties of a solution.

Reversible macromolecular water binding could stabilize intracellular osmolality from fluctuations in extracellular osmolality but only if the plasma membrane were water impermeant. However, the authors experiments show clearly that all the cells that they used have some water permeability, and although they do not say it explicitly in the paper, they show it in their animation. Indeed, all animal cells are permeable to water because phospholipid bilayers are very water permeable (Fettiplace and Haydon, 1980; Mathai et al., 2008).

## Results

To explore how the water permeability of the membrane might be likely to impact the WB hypothesis, we will set up a model cell that incorporates measured values for membrane water permeability. The flux of water across membranes is determined by a very well established physical chemistry relationship that is sometimes called Starling’s equation (Manning and Kay, 2023; Weiss, 1996) for the osmotic volume flux of water is:

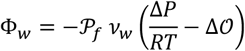

Where 𝒫_*f*_ is the osmotic permeability coefficient, Δ*P* the hydrostatic pressure difference across the membrane, Δ𝒪 the difference in osmolality across the membrane, and 𝒱_*w*_ the molar volume of water.

There is an extensive literature characterizing the 𝒫_*f*_ of lipid bilayer and plasma membrane permeabilities, with the former ranging from 8.3 x 10^-3^ to 1.6 x 10^-3^ cm s^-1^ (Fettiplace and Haydon, 1980; Mathai et al., 2008) while that of human red blood cells, which contain aquaporins, is approximately 2 x 10^-2^ cm s^-1^ (Finkelstein, 1987).

To estimate the impact of the membrane water permeability on a hypothetical WB, we will use the pump-leak model (PLM) that is widely recognized as a mechanism accounting for cell volume stability in the face of extracellular osmotic fluctuations (Fraser and Huang, 2007; Kay, 2017; Rollin et al., 2023; Tosteson and Hoffman, 1960). In this model the action of the Na^+^/K^+^ ATPase (NKA) stabilizes the cell against the osmotic instability induced by impermeant molecules. The PLM drives the cell to a stable volume, with high intracellular K^+^ and low intracellular Na^+^ and Cl^-^. If the extracellular osmolality is changed the cell volume will move to a new steady state where the intracellular osmolality matches the extracellular one.

To simulate the movement of water from putative WBs, we will assume that when the extracellular osmolality is decreased below its steady state value (300 mOsm), water is released at a constant rate. While in the case of an increase in extracellular osmolality, the water is taken up at the same rate by WBs. Fig. 1 shows a simulation for the case of an extracellular osmolality increase (330 mOsm), with and without the WB. As can be seen the WB serves to buffer the change for only a short time, which is determined by the water permeability and the concentration of the condensates. To exaggerate the effect of the WB, we have assumed that the WB represents 10% of the cell volume. The buffering abates in this case when all water has been released from the WB. After that the intracellular osmolality increases until it matches that of the extracellular medium, with the rate being determined by the osmotic permeability coefficient of the membrane, 𝒫_*f*_. Even with 𝒫_*f*_, 2 x 10^-4^ cm s^-1^, an order of magnitude less than that of the lowest lipid bilayer permeability, the addition of water to the cytoplasm preserves the osmolarity and volume for a short time (t_B_ ∼54s, Fig. 1). A smaller change in osmolarity buys more time; an increase in osmolarity to 315 mOsm can resisted for ∼108s. However, with a more realistic value of 𝒫_*f*_ (**1 x 10**^-3^ **cm s**^-1^) t_B_ is ∼20s, It is difficult to argue that this provides any protection, since the buffer is quickly swamped by the osmotic flux of water.

**Fig. 1.**
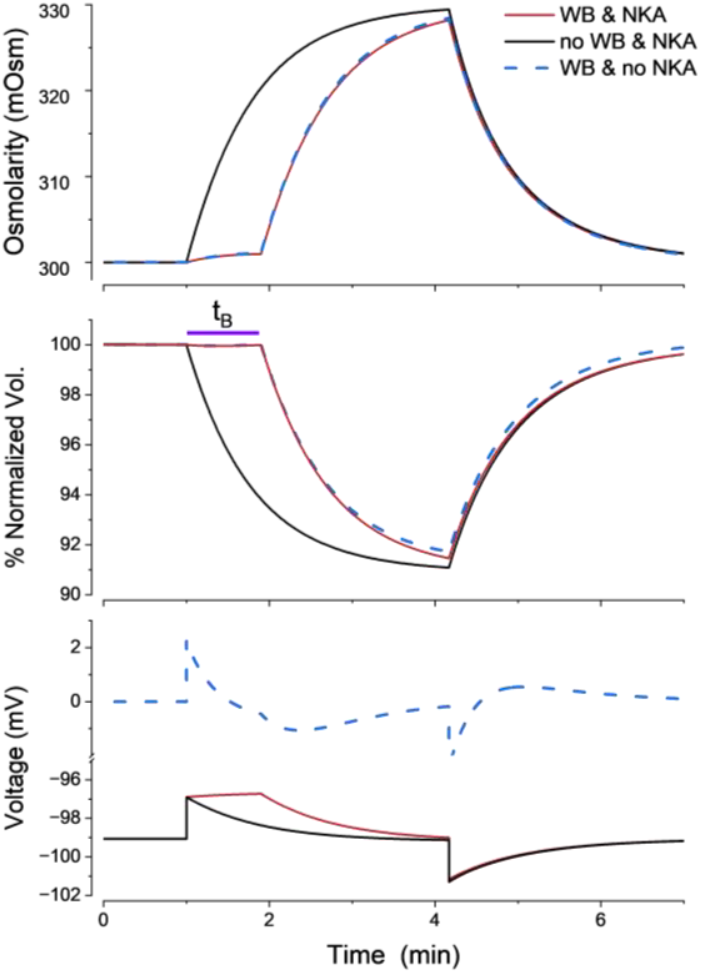
Changes in osmolality, volume and voltage induced by an increase in extracellular osmolality in the presence or absence of a WB. The extracellular solution is initially 147 mM Na^+^, 3 mM K^+^, 130 mM Cl^-^ and 20 mM impermeant anions (X^-^) (300 mOsm) and changed to 330 mOsm at 1 min. The starting radius of the cell is 3.58 *μm*, and the cell contains 2.6×10^-14^ moles of an impermeant molecules with valence -1. The osmotic permeability coefficient, 𝒫_*f*_ of the membrane is 2 x 10^-4^ cm s^-1^. The rate of water release by the WB is 3.3 x 10^-16^ L s^-1^. The cell has Na^+^, K^+^ and Cl^-^ leak conductances and a NKA, all with the same parameters as in Aminzare and Kay (2024). The dashed lines show the responses in the absence of an NKA, with initial radius 3.5 *μm* and 3.58 x10^-14^ moles of impermeant molecules. The system of equations was solved using the stiff ordinary differential equation solver ode15s in MATLAB (R2023b).

To stabilize the intracellular osmolality in the face of changes in extracellular osmolality, the rate of water release from the WB would have to match the rate of water influx through osmosis, which was done in the simulation in Fig. 1. If the rate is not exact, it will lead to swings in osmolarity. To see this consider what would happen if the release from a WB were rapid. An increase in extracellular osmolality will trigger a release of water, which will lower the osmolality below its resting value. The osmolality will then increase to match that of the extracellular solution at a rate set by the water permeability of the membrane (data not shown).

If instead of a hyperosmotic shock, a hypoosmotic shock of 30 mOsm is imposed on the cell, the kinetics of the relaxation to a new steady state is identical (data not shown), however, buffering ceases when the WB is saturated.

The conclusion that a WB is unlikely to be effective in regulating intracellular osmolarity is not dependent on the assumption of a PLM. To demonstrate this we have simulated a cell which is stabilized by the provision of a high concentration of an extracellular impermeant molecule (Aminzare and Kay, 2024) (20mM and valence −1) and has no NKA. For a 30 mOsm increase in osmolarity with the same parameters as in Fig. 1, the t_B_ is the same as that for the case where the NKA is active. Notice that the resting potentials and changes in voltage for the active and passive cases are different (see lower panel Fig, 1).

There are two scenarios, that go beyond the classical PLM, that could potentially salvage the WB hypothesis, namely, the active transport of water, or the development of a turgor pressure, each of which we will consider in turn. If cells could transport water actively, as has been claimed by some (Macaulay et al., 2004), the Starling equation shows that the operation of the water pump would simply be equivalent to an applied transmembrane pressure. In the case of a hyperosmotic shock, the activation of active water transport into the cell stabilizes the volume and hold the intracellular osmolality at a lower value than the outside (data not shown). However, if this were so, the action of a WB would be unnecessary. Evidence has been presented for the active transport of water (Zeuthen and Macaulay, 2013) however, this has been disputed by others (Boron and Boulpaep, 2016; Lapointe et al., 2002; Mollajew et al., 2010).

The influx of water driven by an osmolality difference will lead to the development of tension in the membrane that could limit the expansion of the cell (Rangamani, 2022). If the tension is sufficiently high, the pressure developed could counteract the osmotic pressure and preserve a steady state. In this case it is possible for the osmolality of the intracellular solution to be greater than that of the extracellular solution in the steady state. As an example, if the osmolality of the extracellular solution is dropped by 30 mOsm a hydrostatic pressure of 0.75 atm (calculated from the van ‘t Hoff equation) will develop across the membrane. Using Laplace’s law, one can calculate the membrane tension required to sustain this pressure, which would be ∼ 180 mN/m for a spherical cell with a radius of 5 μm. This is greater than the highest membrane tensions that have been encountered (∼0.3 mN/m) (Kozlov and Chernomordik, 2015). Although higher local and transient increases in tension may occur (Lewis and Grandl, 2015; Shi et al., 2018). In these calculations the membrane is assumed to be smooth, however if the membrane is deformed locally by the cytoskeleton, this could reduce the tension. If the cell has a radius of 5 μm, and the membrane has regular, convex hemi-spherical protrusions with radii of 50 nm, tiling the surface, the tension would be 100 times less than that of a smooth membrane. Notice, if an elevated pressure can be sustained, this alone could preserve an osmotic gradient, without the need for a WB.

In the case of a hyperosmotic shock, the cell would shrink, but there are to the best of our knowledge, no known mechanical mechanisms that could develop a negative pressure to oppose the gradient in osmolality.

## Discussion

Watson *et al*.*’s* hypothesis is ingenious and appears plausible at first glance; however, when incorporated into cell models, which allow the osmotic flux of water, it fails to maintain intracellular osmolality in response to extracellular osmolality changes for more than a minute, even assuming an osmotic permeability ten times lower than typical values. A water buffer could preserve the intracellular osmolality of blood cells in transit through the vasa recta (∼10s) in the kidney (Pallone et al., 1990), which occurs approximately every 5 min for a blood cell (Burton, 2000). However, water buffering would be ineffective in countering changes over a few minutes in duration as might occur during hypo- or hypernatremia (Sterns, 2015).

Watson *et al’s* paper does not consider cell volume regulation, however, if the osmolality of the extracellular saline is changed either the cell volume or the transmembrane pressure must change, or some admixture of the two, because the membrane is permeable to water. Since the plasma membrane is the primary site for water fluxes, one cannot avoid considering changes in volume and pressure, when attempting to make sense of the regulation of the intracellular osmolality. If the WB hypothesis were true, a change in extracellular osmolality should not lead to an immediate change in cell volume, but a volume change should ensue when the water buffer is overwhelmed. If there is no water buffering, there should be a rapid change in volume.

The authors do not consider the PLM, however since it provides the best account of cell volume and osmolality regulation, it is difficult to avoid. As we have shown above, even if the cell is passively stabilized, a WB would be quickly overwhelmed by transmembrane water fluxes. In viewing cellular water fluxes, one cannot sidestep adopting a model. If one assumes that the cell is a closed membrane with a constant volume, if a hypotonic solution bathes the cell, osmosis will drive water in and increase the hydrostatic pressure. If one assumes that the that the plasma membrane is pliant, a decrease in osmolality will lead to the influx of water, and the expansion of volume until the cytoplasmic osmolality matches that outside.

The authors say, “Given the essential roles, high abundance and very large number of different IDR (intrinsically disordered regions)-containing proteins in cells it is impossible to formally prove that rapid changes in the global level of biomolecular condensates enable cells to accommodate acute physiological fluctuations in temperature, osmolality and, presumably, hydrostatic pressure.” However, as we have shown, we are not without means to ward off flawed hypotheses; it is here where theory comes into its own and allows one to assess the likelihood of a hypothesis.

What may, in part, have led the authors to misconstrue the regulation of cytoplasmic osmolality is shown in their Conclusion, where they say “… changes in condensation occur to minimize the perturbation to thermodynamic equilibrium of the system as a whole.” A live cell is not at thermodynamic equilibrium, since it needs to consume energy to stay alive, however, ion concentrations and the cytoplasmic osmolality may be at steady states, but these are nonequilibrium states. Halting all energetic processes lets the cell relax into a thermodynamic equilibrium, which is just another way of saying that it is dead. One of the key conclusions of years of work on cellular physiology, is that although ion concentrations may appear static, they exist in a nonequilibrium state that is driven by continuous energy dissipation (Weiss, 1996; Yang et al., 2021).

The authors’ claim in their Conclusion “… that biomolecular condensation is the primary buffer of intracellular water potential” is incorrect, since they have ignored the plasma membrane, which is the primary site of water fluxes into and out of cells. As we have argued there is a considerable body of knowledge on the osmotic permeability of plasma membranes, which is undergirded by a rigorous theoretical foundation that allows one to predict how WBs are likely to operate in the context of a water permeable membrane. All of this shows that WBs are unlikely to provide protection against changes in extracellular osmolality.

On the experimental side, the authors do not show anything that directly confirms their hypothesis within a live cell. What would be required is that after, say an increase in extracellular osmolality by 30 mOsm, they show that the intracellular osmolality increases rapidly and then is restored quickly to its control value of ∼ 290 mOsm.

While such experiments currently may not be feasible, there have been attempts to develop genetically encoded osmosensors (Kleist et al., 2022), but it will take time to refine and optimize these sensors. It is also worth noting that a nanoscale fluorescent sensor has been developed to sense the water activity within the air spaces in plant leaves (Jain et al., 2021). However, mounting an investigation of a hypothesis that is unlikely to literally hold much water may be quixotic.

Although the authors’ hypothesis may be unlikely, their work could serve to call attention to what is a very important question, the contribution of impermeant molecules to the cytoplasmic osmolality. If the intracellular osmolality is ∼ 290 mOsm, about 100 mOsm is contributed by membrane impermeant molecules (Kay, 2017), a significant question is what fraction of this can be attributed to macromolecules and what to small metabolites (Burton, 1983; Model et al., 2023)?

In conclusion, osmosis is likely to be the primary driver of water transport in cells, so that even if there are cytoplasmic water buffers, a transmembrane difference in osmolality will drive a water flux that will override any WBs.

## Acknowledgements

The authors thank John O’Neill, Emmanuel Derivery, Rachel Edgar, and Joseph Watson, for engaging in an extended email discussion of their paper, Sandipan Chowdhury, Rumiana Dimova, Gerald Manning, Sean Megason, and Aylin Rodan for helpful comments and suggestions on an early version of this paper. The authors are supported by NSF grant 2037828, and ZA by a grant from the Simons Foundation MPS–TSM– 00008005.

## Author contributions

ARK: conceptualization, formal analysis, investigation, writing – original draft, and software.

ZA: formal analysis, software, and writing – review and editing

## Data availability

The MATLAB code will be made available on request.

